# Multidimensional components of impulsivity during early adolescence: Relationships with brain networks and future substance-use in the Adolescent Brian and Cognitive Development (ABCD) study

**DOI:** 10.1101/2025.06.06.658318

**Authors:** Annie Cheng, Steven J. Riley, Robert J. Kohler, Feza Anaise Umutoni, Marc N. Potenza, Sarah D. Lichenstein, Avram J. Holmes, Danilo Bzdok, Sarah W. Yip

**Affiliations:** Yale School of Medicine, Department of Psychiatry, New Haven, CT, USA; Yale University, Wu Tsai Institute and School of Medicine, Departments of Child Study and Neuroscience, New Haven, CT, USA; Connecticut Mental Health Center, New Haven, CT, USA; Connecticut Council on Problem Gambling, Wethersfield, CT, USA; Rutgers University, Department of Psychiatry, Brain Health Institute, Piscataway, NJ, USA; Mila, Montreal; Department of Biomedical Engineering, BIC, MNI, McGill University, Montreal, QC, Canada; Child Study Center, Yale School of Medicine, New Haven, CT, USA

**Keywords:** substance-use risk, connectivity, impulsivity, delay discounting, inhibitory control, positive, negative urgency

## Abstract

Impulsivity is a multifaceted construct that typically increases during adolescence and is implicated in risk for substance use disorders that develop later in life. Here, we take a multivariate approach to identify latent dimensions of impulsivity, broadly defined, among youth enrolled in the Adolescent Brain and Cognitive Development (ABCD) study and explore associations with individual differences in demographics, substance-use initiation and canonical resting state networks (N=11,872, ages ~9-10). Using principal component analysis, we identified eight latent impulsivity dimensions, the top three of which together accounted for the majority of the variance across all impulsivity assessments. The first principal component (PC1) was a general impulsivity factor that mapped onto all impulsivity-related assessments. PC2 mapped onto a ‘mixed’ impulsivity style related to both poorer, less attentive performance on the SST and decreased delay discounting. PC3 linked externalizing behaviors across multiple measures with indices of delay discounting, making delay discounting the only impulsivity-related assessment to load on all three of the top PCs. Multiple impulsivity PCs were significantly associated with subsequent initiation of alcohol and cannabis use. Finally, we found both cross-sectional and longitudinal associations between the PCs and functional connectivity between and within frontoparietal, cingulo-opercular, and default mode networks. These data provide a critical empirical baseline for how facets of impulsivity covary in early adolescence which may be tracked through future waves of ABCD data, enabling longitudinal elucidation of how dimensions of impulsivity interact with other individual and environmental factors to influence risk for substance use later in life.

## Introduction

Impulsivity encompasses a diverse set of interrelated behaviors, including acting without thinking, taking excessive risks, and struggling with delayed gratification [1–6]. Impulsivity typically increases during adolescence as part of normative development. This development, along with co-occurring neurodevelopmental changes, subserves increases in risk-taking behaviors during this time [7, 8]. Risk-taking behaviors during adolescence manifest in diverse ways but, in some individuals, include experimentation with alcohol and drugs-of-abuse. Thus, as earlier ages of initiation of substance-use are associated with significantly increased risk of later development of a substance-use disorder (SUD), adolescence is a critical time of increased vulnerability for addictions and associated negative sequalae [7].

While associations between heightened impulsivity and substance-use and other risk behaviors in youth are well-established, few studies have systematically assessed different facets of impulsivity in a large cohort of youth, in large part due to limited population-level data. The ongoing Adolescent Brain and Cognitive Development (ABCD) study [9, 10] for the first time enables such assessment of impulsivity’s multidimensional structure in a large developmental cohort recruited from diverse sites with harmonized assessments. Here, we use principal component analysis (PCA) to identify latent dimensions of impulsivity, broadly defined, using a wide collection of impulsivity-related assessments from the first wave of ABCD data collection. In doing so, we aim to provide an empirical baseline via which impulsivity may be tracked in later waves, to characterize associations with subsequent substance-use initiation, and to explore associations with individual differences in patterns of brain functional connectivity.

Multiple well-established measures of different aspects of impulsivity exist; however, little is known about the precise relationships between these interrelated, yet partially distinct, impulsivity measures in youth. ABCD includes a diverse collection of commonly administered self- and observer-(e.g., caregiver) reports and behavioral measures related to impulsivity. Behavioral tasks, such as the stop signal task [SST; 11] and delay discounting task [12], measure response and choice impulsivity, respectively [13]. The Behavioral Inhibition/Activation System (BIS/BAS) scales [14] measure avoidance-related and approach-related impulsivity tendencies, respectively, and the Child Behavior Checklist [CBCL; 15] assesses, among other domains, externalizing problems theoretically related to impulsivity. The UPPS-P Impulsive Behavioral Scale [16] measures five facets of impulsivity including sensation-seeking, perseverance, premeditation, and positive and negative urgency. Here, we leverage the unprecedented ABCD dataset to employ a multivariate analysis approach to investigate the relationships between these impulsivity-related measures.

Prior work indicates that dimensions of impulsivity may be differentially associated with specific adolescent risk behaviors, and that the strengths of these associations may further differ as a function of both clinical presentations and basic individual difference factors. For example, a recent literature review [17] found that non-planning and affect-based impulsivity measures—but not motor and choice impulsivity measures—were linked to an increased likelihood of substance-use initiation, whereas increases across all four domains were observed among youth with conduct disorder (CD) [18]. Moreover, demographics and familial factors can moderate the relationship between impulsivity and outcomes. For example, recent work indicates that sex significantly moderates the relationship between impulsivity and negative consequences due to cannabis use in young adults [19]. Given the interactions between various impulsivity dimensions, substance-use, and demographic factors, as evidenced by previous studies, we here consider these core individual difference factors in relation to identified latent dimensions of impulsivity.

Furthermore, as individual differences in neurodevelopment are posited to be central to individual differences in risk-taking behaviors, we further tested for associations between impulsivity dimensions and functional network metrics. While predominant theories of risk behaviors during development typically emphasize a developmental ‘mismatch’ between ‘top-down’ prefrontal cortical inhibitory control regions and ‘bottom-up’ subcortical reward regions, some recent data challenge this assumption [20–22]. In addition, insights from connectivity-based analysis approaches indicate that normative maturation involves complex network-level changes, rather than simple regional development [23–26]. To test the relevance of individual differences in large-scale neural networks in conferring individual differences in impulsivity in youth, we therefore focused on three ‘canonical’ neural networks that, together, are theorized to be critical to basic mechanisms such as attention shifting and resource allocation—namely the frontoparietal, default mode, and cingulo-opercular networks [27]. These three networks not only subserve important mental processes, including working memory, self-referential mental processes, and the detection of motivationally salient information, but they are also suggested to function in an interconnected manner wherein alterations in engagement and/or disengagement of one of these three networks may subserve cognitive dysfunction and risk for psychiatric disorders including SUDs [28–31]. A better understanding of how functional connections within and between these networks are associated with facets of impulsivity during adolescence (as well as the longitudinal trends of these associations) may inform neurodevelopmental models of risk for impulsivity-related psychopathologies.

To achieve the above aims, we apply principal component analysis to impulsivity-related assessments included in ABCD and characterize associations between identified latent components and individual difference factors including future substance-use initiation and brain network properties. Our aim was to disentangle the different facets of impulsivity and to elucidate the complex relationships among these factors while also assessing the status of some outstanding questions related to impulsivity measures in this cohort. We predicted that PCA would identify multiple dimensions of impulsivity, which would differentially map onto individual difference factors. Specifically given the tight association between impulsivity and substance use among adolescents, we predicted that multiple dimensions of impulsivity would be significantly associated with measures of alcohol- and cannabis-use initiation. Finally, we also predicted that the impulsivity dimensions identified at baseline would be associated with connectivity within and between frontoparietal, cingulo-opercular, and default mode networks, both cross-sectionally and longitudinally.

## Methods

An overview of the sample, measures, and methods used in this study can be found in Figure 1.

**Figure 1.**
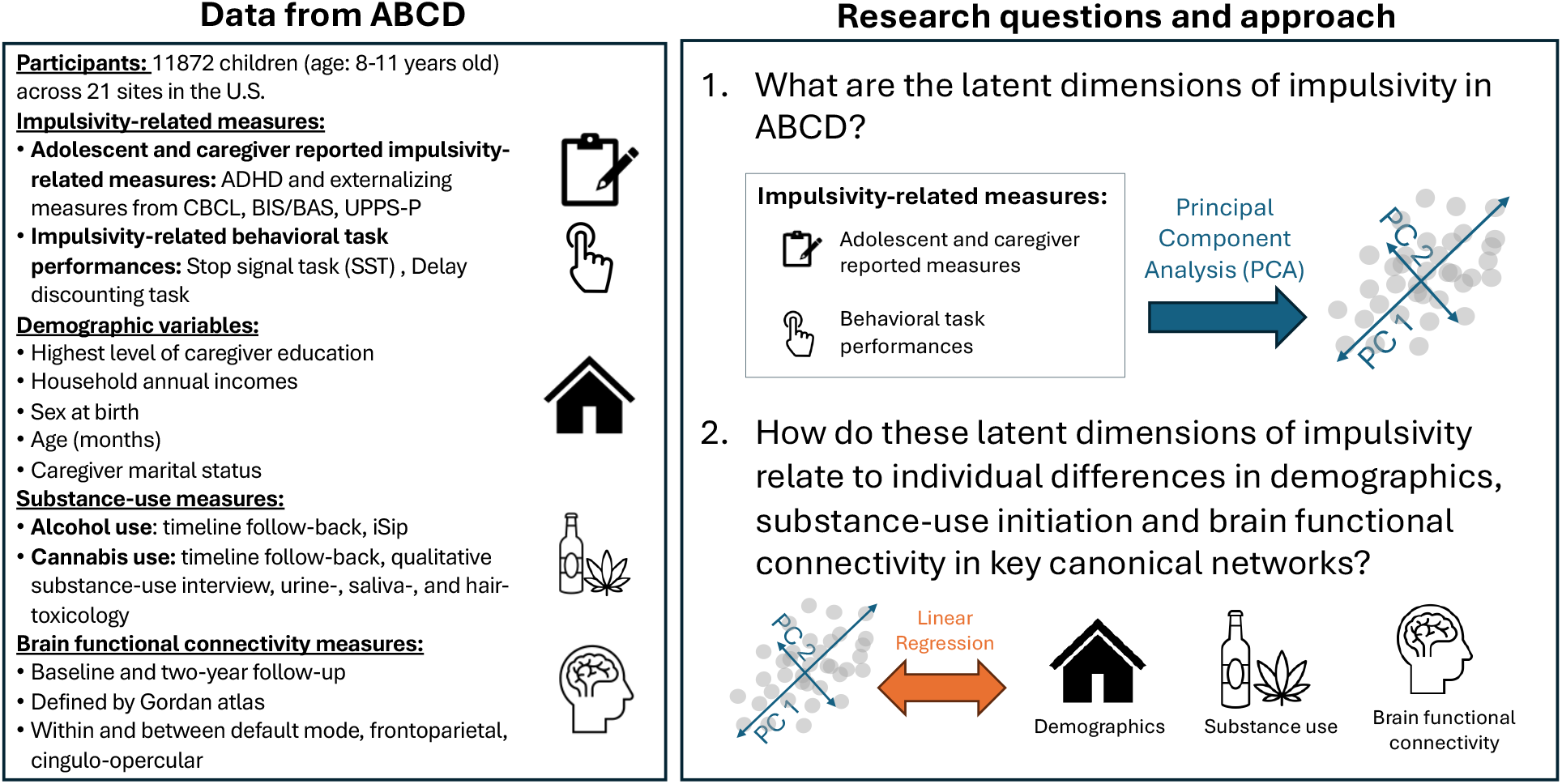
An overview of the data selected from the Adolescent Brain and Cognitive Development (ABCD) study, research questions and data analytic approach. We applied principal component analysis (PCA) to a range of impulsivity-related measures, broadly defined, to identify latent dimensions of impulsivity within the ABCD data set. We then examined the relationships between the top three identified impulsivity PCs and variables related to basic individual difference factors, substance use, and functional connectivity among well-established canonical networks via linear regression analyses.

### Adolescent Brain and Cognitive Development (ABCD) study

Data were obtained from the ABCD study [9] release 4.0 through the National Data Archive in adherence to the stipulations outlined in the data use agreement (NIMH Data Use Agreement #13967). The ABCD study encompasses a sample of 11,872 children, with participants first assessed between the ages and 11. Demographic characteristics of this sample are shown in Table S1.

### Impulsivity-related assessments

Here, we take a broad approach to impulsivity, selecting multiple measures in ABCD with high theoretical relevance to impulsivity and to impulsivity-related disorders. At the behavioral level, this included performance data from the SST and the delay discounting task. The SST [11] is a well-validated task for assessing inhibitory control through a series of GO and STOP trials, where participants must quickly respond to GO signals and inhibit their response when a STOP signal appears. Delay discounting measures the preference for sooner, smaller rewards over later, larger rewards [12]. In the ABCD study, this construct is assessed through a computer-based task where participants make forced choices between immediate and future rewards, with the task scored by seven indifference points that estimate the participant’s subjective value of a future reward at delays ranging from six hours to five years.

The adolescent- and caregiver-reported impulsivity-related measures included are from the BIS/BAS, and UPPS-P and CBCL. The BIS/BAS scales [14] comprise a 24-item self-report questionnaire that assesses the behavioral inhibition and activation systems, which regulate responses to aversive and rewarding stimuli and which map onto a BIS subscale and three BAS subscales: Drive, Fun Seeking and Reward-Responsiveness. The UPPS-P is a 59-item self-report questionnaire that assesses five dimensions of impulsivity: positive urgency, negative urgency, premeditation, perseverance, and sensation seeking. All five subscales from the UPPS-P were included in the current analysis. The CBCL is a 113-question caregiver-reported assessment that evaluates a child’s psychological and behavioral functioning using a three-point Likert scale across eight domains and six diagnostic categories. While the CBCL is not a traditional impulsivity assessment per se, we here include age- and sex-normed t-scores from the ADHD, attention, and externalizing domains from this measure, given the high relevance of these constructs to impulsivity.

### Principal component analysis (PCA)

We performed PCA to identify the main axes of shared co-variation across the above described 28 impulsivity-related assessments. PCA simplifies complex data by combining statistically related features into fewer, more manageable elements, i.e., principal components (PCs). Each observation is then recast as a linear sum of these PCs. This process simplifies the data by reducing each point to a small vector of coefficients. This approach is consistent with prior work using PCA to delineate the structure of cognitive measures in ABCD [34]. Additional details on data scaling, dummy coding and dealing with missing data are provided in the Supplemental Materials.

One potential weakness of ‘standard’ PCA is that it does not distinguish between PCs that result from true vs. spurious correlations among variables [35]. In addition, traditional methods for selecting which PCs are relevant, and methods for deciding which variable loadings within each PC are relevant, typically use ad hoc thresholds [36]. To circumvent these limitations, we used the PCATest library [37] in R [38], which tests the overall PCA, the explained variance of each PC, and the strength of each variable’s loading (i.e., its weight or coefficient on the PC) against permutation-based estimates of null distributions. The false discovery rate (FDR) for significance testing of the PCs and loadings was controlled using the Benjamini-Yekutieli procedure [39], with FDR set at q=.05 (i.e., we expect 5% of significant findings to be false positives).

### Associations between latent impulsivity dimensions and substance-use, demographic variables and brain networks

To maximize the number of participants with substance-use for data analysis, we analyzed the alcohol- and cannabis-use variables from data release 5.1, corresponding to data collected through age 13-14. Alcohol-use is assessed with both a timeline follow-back and the alcohol sipping survey ‘iSip’ [40].

Due to the low prevalence of adolescents who report consuming a full drink of alcohol, and the negative effects associated with sipping alcohol during adolescence [40], the iSip inventory is used to measure low-level use behaviors. For the present study, an adolescent is determined to have used alcohol if they reported at least one sip by the 4-year follow-up for non-religious purposes. This includes reports of a sip or drink during the mid-year phone interviews (i.e., 6-month, 18-month, 30-month, 42-month). As of the 5.1 release, approximately half of the cohort had available iSip data for year four. For those without available year four data, alcohol use variables are calculated based on data through the 3-year follow-up, including mid-year interviews. Cannabis-use is assessed with a combination of data from timeline follow-back, qualitative substance-use interviews, and urine-, saliva-, and hair-toxicology (hair toxicology not available for all participants). For the present study, an adolescent is determined to have used cannabis if any of these measures indicated cannabis-use up to and including the year-four follow-up. Demographic variables included: highest level of caregiver education; household annual incomes in ranges of less than $50,000, between $50,000 and $100,000, and greater than $100,000; sex at birth; age in months; site; and caregiver marital status.

Functional connectivity data from baseline and two-year follow-up was obtained from the curated data release of the ABCD Study [41]. Functional connections of interest included those within and between default mode, frontoparietal and cingulo-opercular networks, as defined by the Gordon atlas [42]. We followed the ABCD-recommended image inclusion criteria to exclude neuroimaging data of poor quality, resulting in N=9601 and N=6929 for baseline and two-year follow-up assessments, respectively. Note that not all data for the two-year follow-up is available in release 5.1, hence the smaller sample size.

## Results

### Latent dimensions of impulsivity

As shown in the scree plot in Figure 2, PCA results indicated that the data were correlated above chance (p<.001) but were not from a single latent variable, consistent with the current understanding of impulsivity as a multi-dimensional construct [43]. In typical unitary data structures, a sharp inflection or “elbow” in the scree plot—where the variance explained by each successive PC markedly declines—is used to distinguish meaningful components from noise [35]. In contrast, the present data deviated from this pattern. The scree plot did not display a clear elbow, and the first two PCs accounted for nearly equal proportions of variance (13.0% and 12.4%, respectively), with the first PC explaining far less than the majority of variance. These features suggest that the data do not cohere around a single dominant factor. Given that the overall PCA structure supported our hypothesis of multidimensionality, we next examined the characteristics of individual PCs.

**Figure 2.**
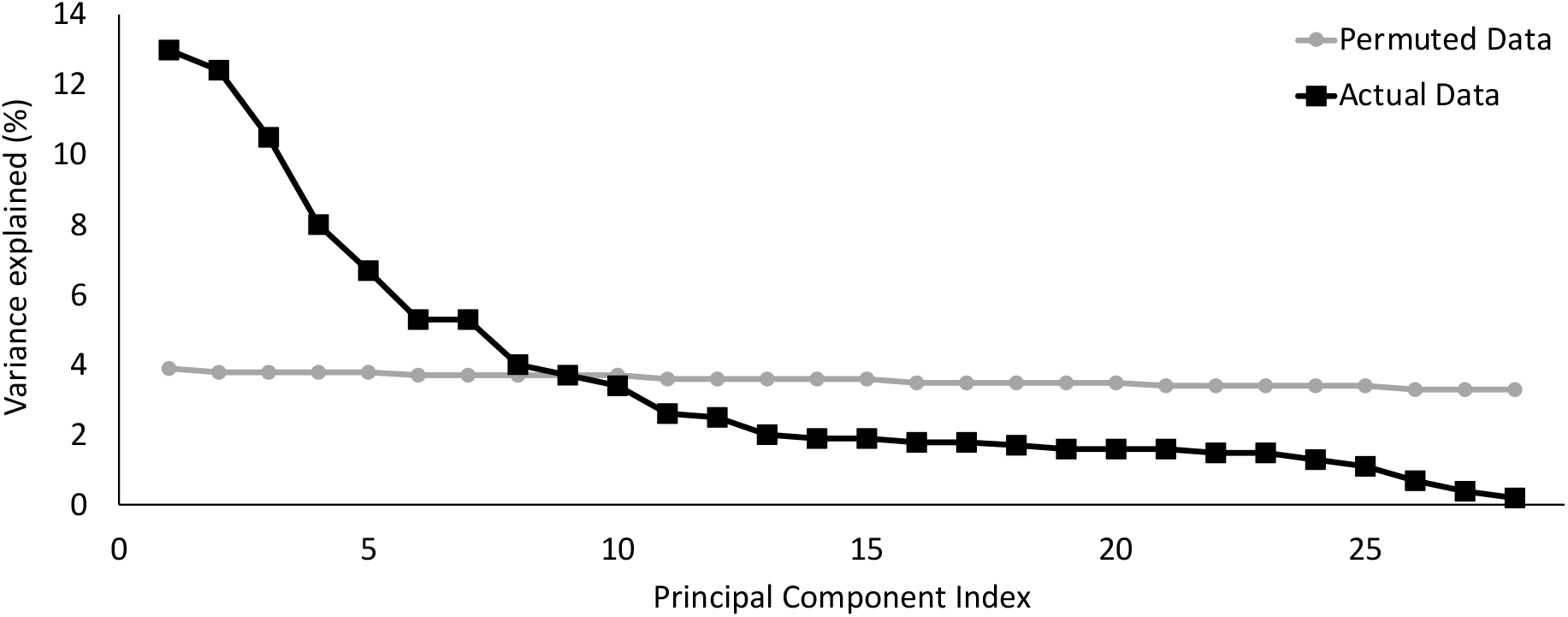
The percent of variance explained by the principal components of the actual data (black line) versus the permutated data (grey line). We did not observe a sharp bend or “knee” at the boundary between PCs that explain a large proportion of variance versus those that do not, indicating that the data does not “hang together” as a unitary construct, consistent with current theories of impulsivity as a multidimensional construct. The first eight PCs emerged as significant as determined using permutation testing.

Eight PCs accounted for statistically significant (q<.05) portions of the total variance as determined by permutation testing and as displayed in the scree plot in Figure 2, together accounting for 65.3% of the variance. Explained variance ranged from 13.0% for PC1 to 4.1% for PC8. Below, we focus on findings from PCs that each accounted for at least 10% of the variance; i.e., PC1, PC2 and PC3 (which together account for 35.9% of the total variance; details of PCs 4-8 in Table S2 and in the SI).

Loadings for the first three PCs are in Figure 3. PC1 included variables from all impulsivity-related assessments (though not all subscales), such that higher scores on this primary axis-of-covariation corresponded to more positive results for variables such as UPPS-P subscale scores and numbers of incorrect STOP trials on the SST and to more negative values for delay discounting indifference points (i.e., increased delay discounting). PC2 was entirely composed of behavioral task variables and revealed a mixed component of impulsivity characterized by poorer SST performance, including both longer RTs and lower accuracy, but decreased delay discounting. PC3 captured shared positive associations between all three CBCL scales (i.e., ADHD, externalizing, attention) and, not surprisingly, externalizing domains of BIS/BAS and UPPS-P scales (i.e., fun-seeking, reward-responsiveness, drive, positive and negative urgency). Interestingly, this component also included multiple delay discounting variables, making delay discounting the only assessment to load significantly on all three of the top PCs.

**Figure 3.**
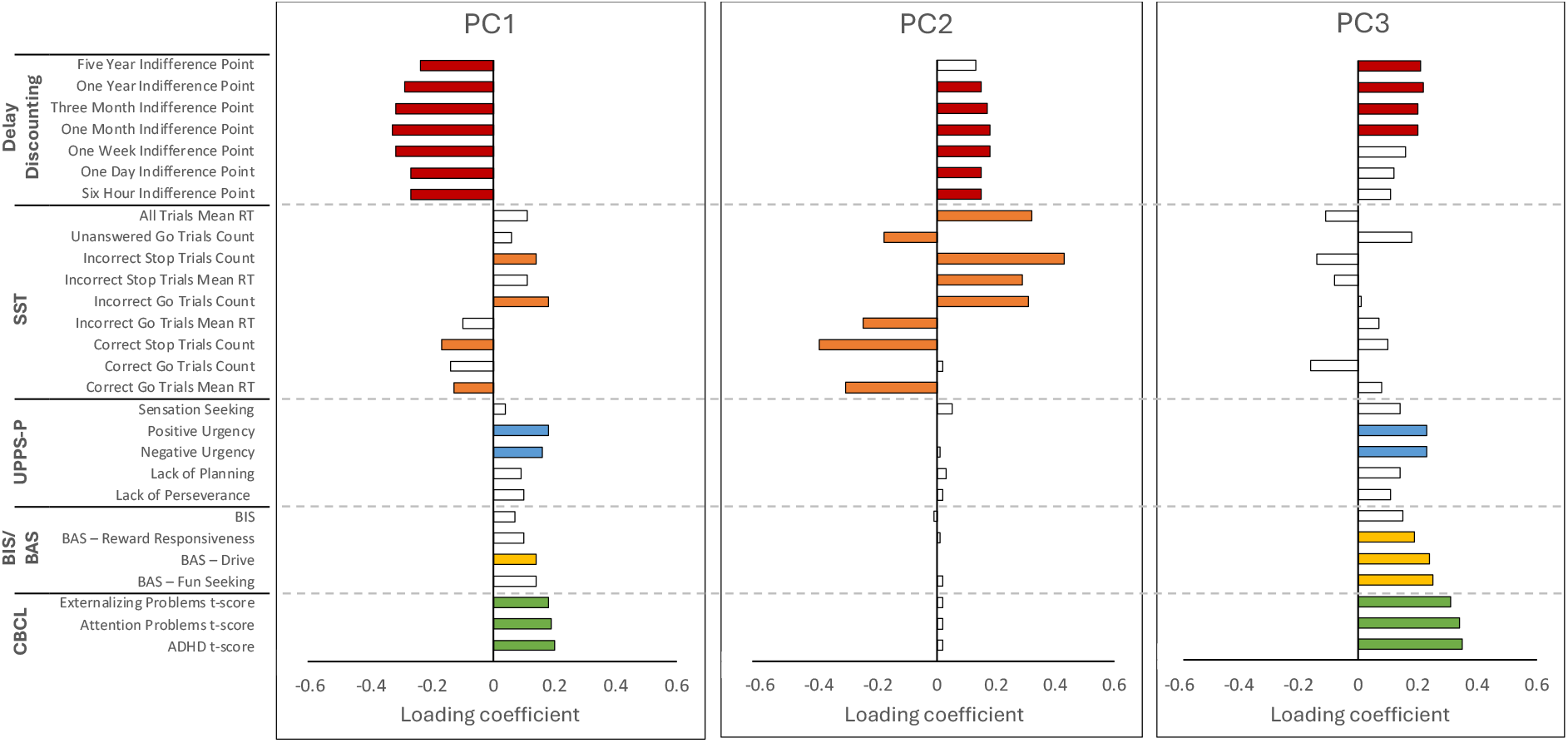
Loading coefficients for impulsivity-related measures onto the top 3 principal components. Colored bars indicate significant loading coefficients, and white bars indicate non-significant loading coefficients. PC1 had significant loadings from at least one variable from each measure; PC2 was entirely composed of behavioral task variables and revealed a dimension-of-covariation ranging from negative loadings for total correct ‘stops’ to positive loadings for total incorrect ‘stops’ on the SST, with delay discounting measures in between these two extremes; PC3 captured shared positive associations between all of the three CBCL scales (i.e., ADHD, externalizing, attention), externalizing domains of BIS/BAS and UPPS scales, and multiple delay discounting variables.

### Relationships with substance-use, demographic variables and brain networks

PC scores for each of the eight significant PCs were calculated for all participants and entered into linear regression to determine relationships with substance-use and demographic variables. There were 288 total comparisons (eight PCs by 36 variables per regression), of which 92 were significantly different from zero (q<.05). Coefficients are listed in Table S3, and significant coefficients for the first three components are shown in Figure 4. Consistent with our hypothesis, the first three PCs were significantly associated with alcohol use initiation. Standardized regression coefficients for alcohol initiation on the first three PCs were .074, −.074, and .097, respectively. PC1 and PC3 were significantly associated with cannabis-use initiation, with standardized regression coefficients of .049 and .094, respectively. Thus, PC1 and PC3 were each positively related to both cannabis and alcohol initiation, while PC2 was negatively related to alcohol initiation. Sex at birth and age at baseline were significantly related to PCs 1-3. We also observed household yearly income (>$50,000/year) and caregiver education (Post-Graduate, >=Bachelor’s) to be negatively associated with PC1. Caregiver marital status was unrelated to any dimension.

**Figure 4.**
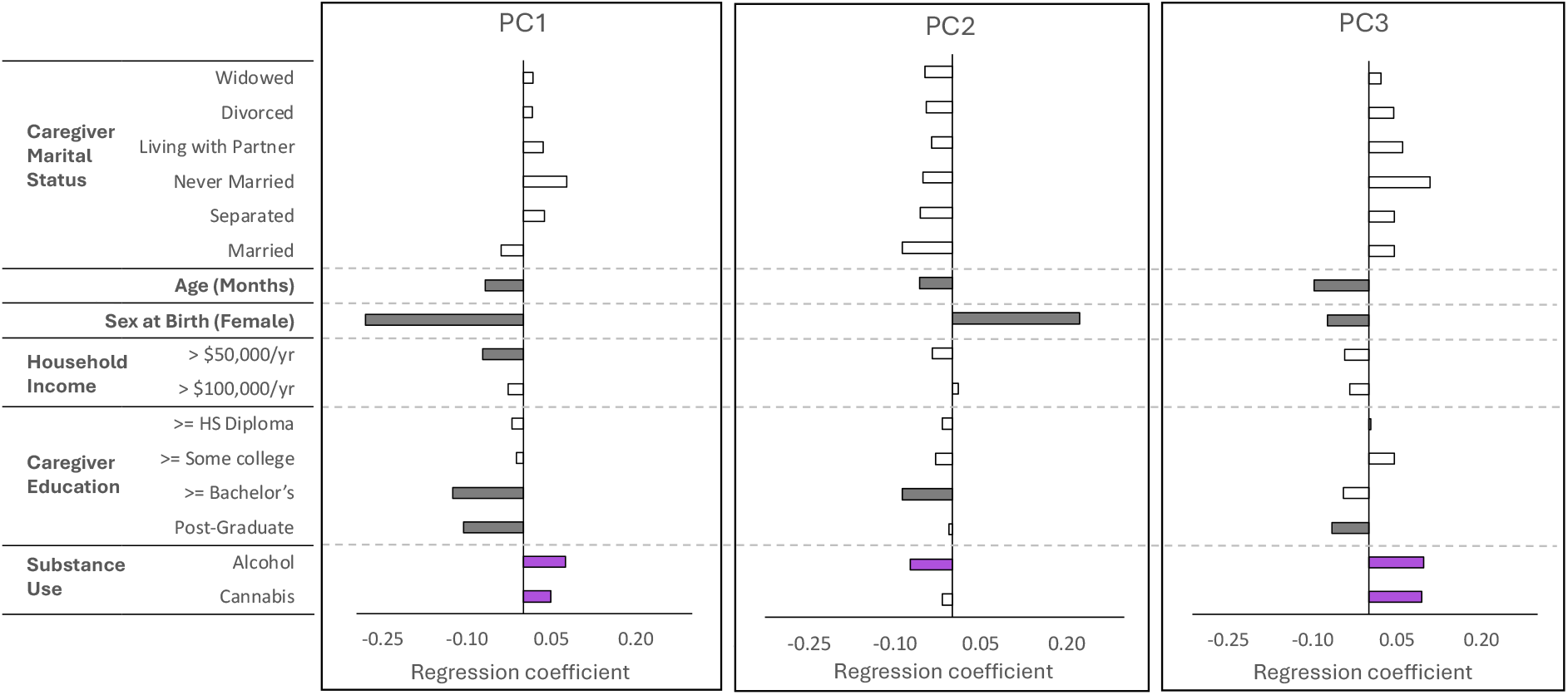
Regression coefficients between principal components and substance use and demographic variables. Colored/darkened bars indicate significant regression coefficients, and white bars indicate non-significant regression coefficients. PC1 and PC3 were each positively and significantly related to both cannabis and alcohol initiation, while PC2 was negatively and significantly related to alcohol initiation, but not cannabis initiation. Sex at birth and age were significantly associated with all three PCs. Household yearly income (>$50,000/year) and caregiver education (Post-Graduate, >=Bachelor’s) were negatively associated with PC1. Caregiver marital status was unrelated to any dimension. The false discovery rate (FDR) for significance testing of the PCs and loadings was controlled using the Benjamini-Yekutieli procedure with FDR set at q=.05

Finally, we tested for an association between the top three PCs and all combinations of connectivity between and within default mode, frontoparietal, and cingulo-opercular networks at baseline and at 2-year follow-up assessments (Bonferroni corrected; Figure 5). PC1 was positively and significantly associated with functional connections between the frontoparietal and cingulo-opercular networks, and between cingulo-opercular and default mode networks, and negatively associated with within-network connections in the frontoparietal, default mode, and cingulo-opercular networks. These relationships were observed both at the baseline and at two-year follow-up assessments, indicating potential stability of these relationships over time. PC2 was associated with connectivity between frontoparietal and default mode network at baseline but not at the two-year follow-up assessment. PC3 shared similar patterns of associations as PC1 at baseline, but the associations were less consistent longitudinally, with only the positive associations between cingulo-opercular and default mode network, and the negative associations within the frontoparietal and default mode networks remaining significant at the 2-year follow-up assessment. Additional results from a partial correlation analysis (controlling for motion during scanning) are included the Supplemental Materials but left primary findings unchanged.

**Figure 5.**
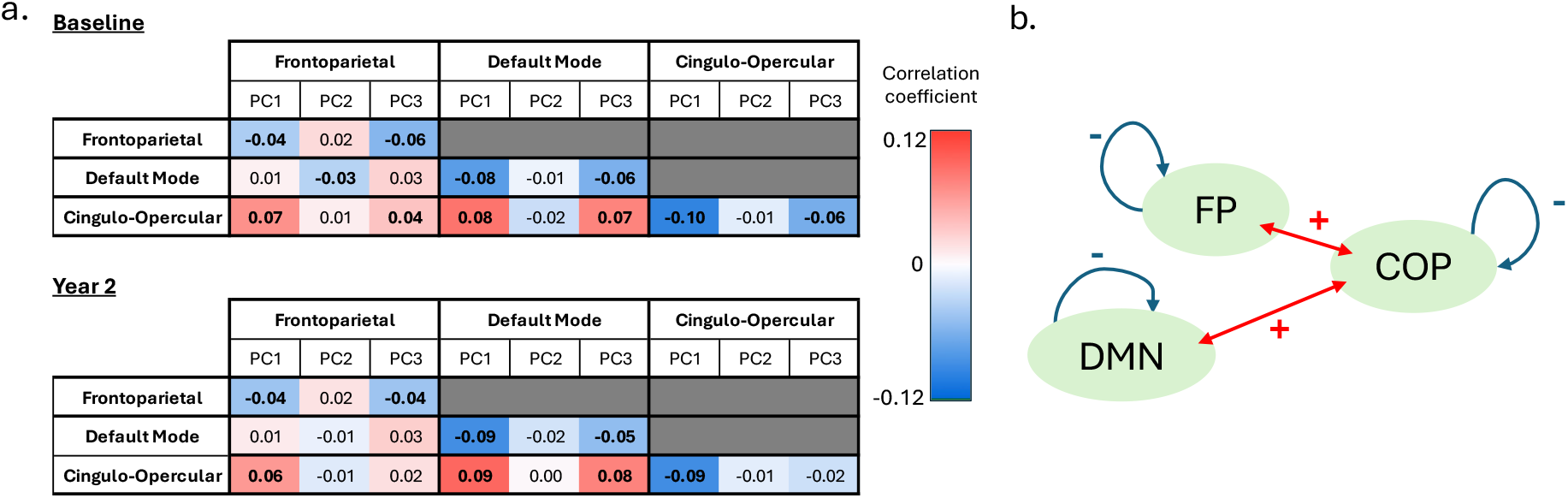
**a**. Correlation coefficients between the top three principal components and all possible combinations of connectivity between and within default mode, frontoparietal, and cingulo-opercular networks at baseline and at 2-year follow-up assessments. Bold font indicates that the coefficient is significant after controlling for false discovery rate. Both at baseline and at two-year follow-up assessments, PC 1 was positively and significantly associated with functional connections between the frontoparietal and cingulo-opercular networks, and between cingulo-opercular and default mode networks, and negatively associated with within-network connections in the frontoparietal, default mode, and cingulo-opercular networks. PC 2 was associated with connectivity between frontoparietal and default mode networks at baseline but not at the two-year follow-up assessment. PC 3 shared similar patterns of associations as PC 1 at baseline, but the associations were less consistent longitudinally. Bonferroni-corrected p<.05. **b**. A schematic representation of the significant associations found between the PCs and the functional connections tested.

## Discussion

This study sought to characterize latent dimensions of impulsivity among youth in the ABCD study, an unparalleled sample of >11,000 young adolescents. It further tested whether identified dimensions map on to individual differences in substance-use initiation (one of the prototypical risk behaviors commonly observed during adolescence), demographic, and brain network variables. We identified eight PCs that accounted for a statistically significant portion of the variance structure across the 28 impulsivity-related variables included in ABCD’s assessment battery, confirming that impulsivity is a multi-dimensional construct. As above, here we focus on findings related to the first three of these components, which together accounted for the majority of variance, while an expanded discussion that includes the remaining five may be found in the SI.

PC1 included variables from all impulsivity-related assessments (though not all subscales). This PC, which may be thought of as a sort of ‘general impulsivity’ dimension, captured common facets of impulsivity, including self-/caregiver-reports related to motivation and urgency that often influence decision-making processes [44, 45] and social interactions [46–48], as well as performance on behavioral tasks. Adolescents with higher scores on PC1 exhibited heightened levels of externalizing behaviors, as indicated by significant positive loadings on the ADHD, attention, and externalizing subscales of the CBCL. Individual differences on in this primary impulsivity dimension were further linked to future patterns of substance-use, as assessed over subsequent waves of ABCD data collection, such that higher scores on PC1 were positively associated with the likelihood of both cannabis- and alcohol-use initiation.

PC2 was composed exclusively of behavioral task variables and reflected a dimension of covariation ranging from negative loadings for total correct stops to positive loadings for total incorrect stops on the SST, with delay discounting measures loading in between these two extremes. Thus, unlike PCs 1 and 3, for which increased loadings may be interpreted as indicative of increased impulsivity, PC2 emerged as a ‘mixed impulsivity’ component, such that higher loadings on this dimension were associated with poorer SST performance and decreased delay discounting (i.e., higher indifference points). Interestingly, PC2 was the only of the top three PCs to emerge as a negative predictor of alcohol-use initiation.

PC3 reflected a common dimension linking elevated scores across all three CBCL scales (ADHD, externalizing, and attention problems) with higher levels of externalizing tendencies on the BIS/BAS and UPPS-P measures (i.e., fun-seeking, reward responsiveness, drive, and urgency). Notably, this component also encompassed several delay discounting indices, distinguishing delay discounting as the only assessment domain to show significant loadings across all the first three principal components. However, somewhat paradoxically, participants with higher scores on PC3 displayed *increased* expression of externalizing behaviors, broadly defined, yet exhibited an increased tolerance for reward-delay—particularly over longer time intervals (i.e., months to years, not days)—indicating *decreased* delay discounting. This seemingly discrepant result raises an important consideration: prior studies have suggested that, in younger adolescents, delay discounting may reflect planning ability more than impulsivity, due to decreased orienting toward the future in younger versus older adolescents [49, 50]. Thus, it is possible that this counterintuitive finding may be driven by decreased orienting or planning over longer time intervals (as shorter delay discounting windows did not load on this PC). In line with this interpretation, PC3, as with PC1, was a positive predictor of both alcohol- and cannabis-use, broadly consistent with general conceptions of impulsivity as a risk factor for earlier ages of substance-use initiation.

Among the demographic variables examined, sex at birth and age in months showed the most widespread associations across the components, underscoring their broad relevance to individual differences in this dataset. Socioeconomic indicators and parental features, including higher household income and caregiver education, were further negatively associated with PC1, suggesting this dimension may capture variability linked to socioeconomic disadvantage. In contrast, caregiver marital status did not relate to any of the identified components. Interpretation of basic demographic variables within this context is challenging, in part due to the complexity of the ABCD dataset itself. Further analysis, powered to assess interactions between variables—i.e., the interconnected nature of social identities—is needed to contextualize these and other findings [51]. For example, our analyses revealed effects of site on PCs (see SI), consistent with prior PCA work focused on cognition in ABCD [34]. However, given the complexity of interpreting site effects, race/ethnicity effects and effects related to parents’ country of origin within ABCD as solitary variables—and without careful consideration of interactions across and between other core features [52]—we believe that interpretation of such effects is as beyond the scope of the present investigation. For example, site may be related to several confounding variables that might affect impulsivity, such as access to nature [53] and ozone exposure [54]. In addition, as our prior work indicates significant effects of social determinants of inequity in the ABCD sample [55], we here did not explore effects related to race and ethnicity in the current manuscript as we do not believe we would be sufficiently powered to explore critical issues of intersectionality pertinent to appropriate interpretation of these variables [51].

To ground the identified latent impulsivity dimensions in a neurodevelopmental framework, we further explored associations between the top three PCs and functional connectivity within and between three large-scale neural networks, i.e., default mode, frontoparietal, and cingulo-opercular networks, previously linked to transdiagnostic features of impulsivity in a smaller neurodevelopmental cohort [56]. We found that PC1 was positively associated with increased connectivity between the cingulo-opercular and frontoparietal networks, as well as between the cingulo-opercular and default mode networks, but not between the frontoparietal and default mode networks. This pattern aligns with prior evidence indicating that the cingulo-opercular network plays a regulatory role in modulating engagement of the other two networks—a function that may be particularly relevant to impulsivity-related phenotypes [57]. We also found the associations between PC1 and these functional connections to be relatively stable longitudinally, highlighting the potential relevance of these connections for the neurodevelopment of impulsivity-related psychopathologies during adolescence. In contrast, associations between connectivity patterns and PCs 2 and 3 exhibited less consistency over time, raising the possibility that the role of these networks in conferring individual differences in impulsivity may shift over the course of development.

### Strengths, limitations, and future directions

A strength of this paper is the size of the sample and its inclusion of diverse youth recruited from multiple sites across the United States. The sample size and the increased power it provides for significance testing allowed for detection of effects that may have been missed in smaller samples that can only detect large effect sizes. There are several limitations that should be mentioned. First, not all data were collected at the same assessment timepoint. For example, delay discounting data were collected during the one-year follow-up, while the SST data were collected at the baseline assessment. Thus, temporal differences in data collection may mask or overrepresent certain effects of age. Second, our analyses primarily focused on the baseline and 1-year variables. Similar analyses of subsequent data releases of ABCD will be required to determine the temporal stability of our findings over development. Whether a general impulsivity PC1 will continue to exist in future releases is an important question, since previous impulsivity measure PCAs have lacked such a component. In addition, future studies are also needed to determine which, if any, of the identified dimensions can reliably predict developmental or psychiatric outcomes beyond initiation of alcohol- and cannabis-use. Another important direction will be considering the relationship between identified PCs from this study and other neurodevelopmental variables, such as brain morphology measures. In summary, this analysis provides a compact set of latent impulsivity dimensions and correlated substance-use, demographic measures and functional connectivity relationships among major brain networks in a large, nationally representative sample of US youths. We demonstrate that the impulsivity measures from the ABCD data set capture several independent constructs. This analysis provides an empirical baseline for future studies of impulsivity in the ABCD dataset, which may leverage the multiyear design of ABCD to examine temporal dynamics in the patterns found in this analysis.

## Authorship

Study design, interpretation of results and manuscript editing was completed by all authors. Primary analyses were conducted by Dr. Riley, in collaboration with Drs. Yip and Cheng. Drs. Riley, Cheng and Kohler contributed to the first round of manuscript writing, with subsequent editing consultation with Dr. Yip.

### Funding

This work was supported by NIDA grant R01DA053301 (MPIs Yip, Bzdok) by CIHR grant 470425 (MPIs Yip, Bzdok), and T32DA022975 (SR). Data used in the preparation of this article were obtained from the Adolescent Brain Cognitive Development (ABCD) Study (https://abcdstudy.org), held in the NIMH Data Archive (NDA). This is a multisite, longitudinal study designed to recruit more than 10,000 children age 9–10 and follow them over 10 years into early adulthood. The ABCD Study® is supported by the National Institutes of Health and additional federal partners under award numbers U01DA041048, U01DA050989, U01DA051016, U01DA041022, U01DA051018, U01DA051037, U01DA050987, U01DA041174, U01DA041106, U01DA041117, U01DA041028, U01DA041134, U01DA050988, U01DA051039, U01DA041156, U01DA041025, U01DA041120, U01DA051038, U01DA041148, U01DA041093, U01DA041089, U24DA041123, U24DA041147. A full list of supporters is available at https://abcdstudy.org/federal-partners.html. A listing of participating sites and a complete listing of the study investigators can be found at https://abcdstudy.org/consortium_members/. ABCD consortium investigators designed and implemented the study and/or provided data but did not necessarily participate in the analysis or writing of this report. This manuscript reflects the views of the authors and may not reflect the opinions or views of the NIH or ABCD consortium investigators. The ABCD data used in this report came from NDA Release 4.0 (DOI: 10.15154/1519007). DOIs can be found at https://dx.doi.org/10.15154/1519007.

### Statement

During the preparation of this work the authors used ChatGPT to improve readability and language. After using this tool/service, the authors reviewed and edited the content as needed and take full responsibility for the content of the publication.

## References

1. Cyders, M.A. and G.T. Smith, Emotion-based dispositions to rash action: positive and negative urgency. Psychol Bull, 2008. 134(6): p. 807–28.

2. Bechara, A., Decision making, impulse control and loss of willpower to resist drugs: a neurocognitive perspective. Nat Neurosci, 2005. 8(11): p. 1458–63.

3. Dickman, S.J., Functional and dysfunctional impulsivity: personality and cognitive correlates. J Pers Soc Psychol, 1990. 58(1): p. 95–102.

4. Franken, I.H.A. and P. Muris, Gray’s impulsivity dimension: A distinction between Reward Sensitivity versus Rash Impulsiveness. Personality and individual differences, 2006. 40(7): p. 1337–1347.

5. Kim, S.J., H.J. Kim, and K. Kim, Time Perspectives and Delay of Gratification - The Role of Psychological Distance Toward the Future and Perceived Possibility of Getting a Future Reward. Psychology Research and Behavior Management, 2020. 13: p. 653–663.

6. Moeller, F.G., et al., Psychiatric aspects of impulsivity. Am J Psychiatry, 2001. 158(11): p. 1783–93.

7. Chambers, R.A., J. Taylor, and M. Potenza, Developmental Neurocircuitry of Motivation in Adolescence: A Critical Period of Addiction Vulnerability. American Journal of Psychiatry, 2003. 160(6): p. 1041–1052.

8. Yip, S.W. and M.N. Potenza, Application of Research Domain Criteria to childhood and adolescent impulsive and addictive disorders: Implications for treatment. Clin Psychol Rev, 2018. 64: p. 41–56.

9. Casey, B.J., et al., The Adolescent Brain Cognitive Development (ABCD) study: Imaging acquisition across 21 sites. Dev Cogn Neurosci, 2018. 32: p. 43–54.

10. Garavan, H., et al., Recruiting the ABCD sample: Design considerations and procedures. Dev Cogn Neurosci, 2018. 32: p. 16–22.

11. Logan, G.D., W.B. Cowan, and K.A. Davis, On the ability to inhibit simple and choice reaction time responses: a model and a method. J Exp Psychol Hum Percept Perform, 1984. 10(2): p. 276–91.

12. Odum, A.L., Delay discounting: I’m ak, you’re ak. Journal of the experimental analysis of behavior, 2011. 96(3): p. 427–439.

13. Romer, D., Adolescent risk taking, impulsivity, and brain development: implications for prevention. Dev Psychobiol, 2010. 52(3): p. 263–76.

14. Carver, C.S. and T.L. White, Behavioral inhibition, behavioral activation, and affective responses to impending reward and punishment: the BIS/BAS scales. Journal of personality and social psychology, 1994. 67(2): p. 319.

15. Achenbach, T.M. and C. Edelbrock, Child behavior checklist. Burlington (Vt), 1991. 7: p. 371–392.

16. Whiteside, S.P. and D.R. Lynam, The five factor model and impulsivity: Using a structural model of personality to understand impulsivity. Personality and individual differences, 2001. 30(4): p. 669–689.

17. Verdejo-Garcia, A. and N. Albein-Urios, Impulsivity traits and neurocognitive mechanisms conferring vulnerability to substance use disorders. Neuropharmacology, 2021. 183: p. 108402.

18. Brislin, S.J., et al., Heterogeneity within youth with childhood-onset conduct disorder in the ABCD study. Frontiers in psychiatry, 2021. 12: p. 701199.

19. VanderVeen, J.D., A.R. Hershberger, and M.A. Cyders, UPPS-P model impulsivity and marijuana use behaviors in adolescents: A meta-analysis. Drug Alcohol Depend, 2016. 168: p. 181–190.

20. Crone, E.A. and R.E. Dahl, Understanding adolescence as a period of social-affective engagement and goal flexibility. Nat Rev Neurosci, 2012. 13(9): p. 636–50.

21. Yip, S.W. and M.N. Potenza, Application of Research Domain Criteria to childhood and adolescent impulsive and addictive disorders: Implications for treatment. Clin Psychol Rev, 2016.

22. Mills, K.L., et al., The developmental mismatch in structural brain maturation during adolescence. Dev Neurosci, 2014. 36(3-4): p. 147–60.

23. Scheinost, D., et al., Sex differences in normal age trajectories of functional brain networks. Human Brain Mapping, 2015. 36(4): p. 1524–35.

24. Farrant, K. and L.Q. Uddin, Asymmetric development of dorsal and ventral attention networks in the human brain. Dev Cogn Neurosci, 2015. 12: p. 165–74.

25. Cai, L., Q. Dong, and H. Niu, The development of functional network organization in early childhood and early adolescence: A resting-state fNIRS study. Developmental Cognitive Neuroscience, 2018. 30: p. 223–235.

26. Di Martino, A., et al., Unraveling the miswired connectome: a developmental perspective. Neuron, 2014. 83(6): p. 1335–53.

27. Menon, V., Large-scale brain networks and psychopathology: a unifying triple network model. Trends Cogn Sci, 2011. 15(10): p. 483–506.

28. Zhang, R. and N.D. Volkow, Brain default-mode network dysfunction in addiction. NeuroImage, 2019. 200: p. 313–331.

29. Cushnie, A.K., W. Tang, and S.R. Heilbronner, Connecting Circuits with Networks in Addiction Neuroscience: A Salience Network Perspective. Int J Mol Sci, 2023. 24(10).

30. Lacomba-Arnau, E., A. Martínez-Molina, and A. Barrós-Loscertales, Structural Cerebellar and Lateral Frontoparietal Networks are altered in CUD: An SBM Analysis. Addict Biol, 2025. 30(3): p. e70021.

31. Sadaghiani, S. and M. D’Esposito, Functional Characterization of the Cingulo-Opercular Network in the Maintenance of Tonic Alertness. Cereb Cortex, 2015. 25(9): p. 2763–73.

32. Tenenbaum, J.B., V. de Silva, and J.C. Langford, A global geometric framework for nonlinear dimensionality reduction. science, 2000. 290(5500): p. 2319–23.

33. Bzdok, D. and B.T.T. Yeo, Inference in the age of big data: Future perspectives on neuroscience. NeuroImage, 2017. 155: p. 549–564.

34. Thompson, W.K., et al., The structure of cognition in 9 and 10 year-old children and associations with problem behaviors: Findings from the ABCD study’s baseline neurocognitive battery. Dev Cogn Neurosci, 2019. 36: p. 100606.

35. Jackson, D.A., Stopping rules in principal components analysis: a comparison of heuristical and statistical approaches. Ecology, 1993. 74(8): p. 2204–2214.

36. Vasco, M., Permutation tests to estimate significances on Principal Components Analysis. Computational Ecology and Software, 2012. 2(2): p. 103.

37. Camargo, A., PCAtest: testing the statistical significance of Principal Component Analysis in R. PeerJ, 2022. 10: p. e12967.

38. Team, R.C., R: A Language and Environment for Statistical Computing. 2022, R Foundation for Statistical Computing: Vienna, Austria.

39. Benjamini, Y. and D. Yekutieli, False discovery rate-adjusted multiple confidence intervals for selected parameters. Journal of the American Statistical Association, 2005. 100(469): p. 71–81.

40. Jackson, K.M., et al., The prospective association between sipping alcohol by the sixth grade and later substance use. J Stud Alcohol Drugs, 2015. 76(2): p. 212–21.

41. Hagler, D.J., Jr., et al., Image processing and analysis methods for the Adolescent Brain Cognitive Development Study. NeuroImage, 2019. 202: p. 116091.

42. Gordon, E.M., et al., Generation and Evaluation of a Cortical Area Parcellation from Resting-State Correlations. Cereb Cortex, 2016. 26(1): p. 288–303.

43. Bakhshani, N.M., Impulsivity: a predisposition toward risky behaviors. Int J High Risk Behav Addict, 2014. 3(2): p. e20428.

44. Jeske, D., P. Briggs, and L. Coventry, Exploring the relationship between impulsivity and decision-making on mobile devices. Personal and Ubiquitous Computing, 2016. 20: p. 545–557.

45. Canale, N., et al., Trait urgency and gambling problems in young people by age: the mediating role of decision-making processes. Addict Behav, 2015. 46: p. 39–44.

46. Sperry, S.H., et al., Examining the multidimensional structure of impulsivity in daily life. Personality and individual differences, 2016. 94: p. 153–158.

47. Jones, K.A., A. Chryssanthakis, and M.J. Groom, Impulsivity and drinking motives predict problem behaviours relating to alcohol use in university students. Addict Behav, 2014. 39(1): p. 289–96.

48. Billieux, J., et al., The role of urgency and its underlying psychological mechanisms in problematic behaviours. Behav Res Ther, 2010. 48(11): p. 1085–96.

49. Steinberg, L., et al., Age differences in future orientation and delay discounting. Child development, 2009. 80(1): p. 28–44.

50. Kohler, R.J., S.D. Lichenstein, and S.W. Yip, Hyperbolic discounting rates and risk for problematic alcohol use in youth enrolled in the Adolescent Brain and Cognitive Development study. Addict Biol, 2022. 27(2): p. e13160.

51. Dhamala, E., et al., Considering the interconnected nature of social identities in neuroimaging research. Nature neuroscience, 2024.

52. Brown, S.A., et al., Responsible Use of Population Neuroscience Data: Toward Standards of Accountability and Integrity. J Adolesc Health, 2024. 75(5): p. 703–705.

53. Repke, M.A., et al., How does nature exposure make people healthier?: Evidence for the role of impulsivity and expanded space perception. PLoS One, 2018. 13(8): p. e0202246.

54. Burkhardt, J., et al., The effect of pollution on crime: Evidence from data on particulate matter and ozone. Journal of Environmental Economics and Management, 2019. 98: p. 102267.

55. Yip, S.W., et al., Multivariate, Transgenerational Associations of the COVID-19 Pandemic Across Minoritized and Marginalized Communities. JAMA Psychiatry, 2022. 79(4): p. 350–358.

56. Jones, J.S., et al., Testing the triple network model of psychopathology in a transdiagnostic neurodevelopmental cohort. Neuroimage Clinical, 2023. 40: p. 103539.

57. Filbey, F.M., et al., Differential associations of combined vs. isolated cannabis and nicotine on brain resting state networks. Brain Struct Funct, 2018. 223(7): p. 3317–3326.

