## Supplemental Information for "Multidimensional components of impulsivity during early adolescence: Relationships with brain networks and future substance-use in the Adolescent Brian and Cognitive Development (ABCD) study"

Supplemental Materials

*Scaling, missing data and dummy coding*

Since PCA is sensitive to the magnitude and the unit of measurement variables [35], each measure was scaled to have a mean of zero and standard deviation of one (i.e., standardized) prior to computing the PCA.

As missing data is often not missing at random, data imputation was performed to maximize data inclusion and to avoid dropping participants with incomplete data. The maximum number of missing data points for an individual measure was 811, out of the total sample of 11872, or 6.8%, and the average rate of missing data across all measures was <3%. Missing data points for each measure were imputed randomly from the set of all valid observations for that measure. This conservative method, known as non-parametric single-column imputation, a form of hot-deck imputation, produces low bias in missing data simulations, where the typical effect is a slight reduction in the correlation between variables [36].

Associations between the identified PCs and the above variables were tested using linear regressions with the PCs as the dependent variable. Dummy-coded variables included site (reference group: site 1 - Children’s Hospital of Los Angeles), caregiver marriage status (reference group: “Refused to answer”), and binary variables (e.g., sex at birth, initiation of alcohol or cannabis). Caregiver education level and household income were ordinally coded. T-tests were performed to detect which regression coefficients were significantly different from zero. Significance levels in regression coefficient t-tests were controlled using the Benjamini-Yekutieli procedure with FDR set at q=.05. Demographic information about the sample is listed in Table S1.

Figures, tables, and discussion related solely to PC1-PC3 are in the main manuscript. Here we provide the data and discussion for the remaining PCs. Table S2 contains the loading coefficients for all variables and all PCs; Table S3 contains the regression coefficients for all substance use and demographic variables where the dependent variable is the relevant PC value.

*Description of primary findings related to PCs 4-8*

All eight PCs are significantly related to alcohol-use initiation, while only two (viz. PC1 and PC3) are related to cannabis use. Among the remainder of demographic and substance use variables, the most common independent variables related to PCs are age (related to seven PCs), and sex at birth (related to six PCs). Data collected at specific sites is also significantly related to many PCs: data from Medical University of South Carolina is related to six PCs, while data collected at Yale, SRI International, and Florida International are each significantly related to five PCs.

Component 4 (PC 4) highlights the distinction between sensitivity to reward and punishment, as measured by the BIS/BAS, and presence of externalizing behavior, as measured by the CBCL. Participants with high scores on PC 4 have CBCL syndrome scale scores above what would be expected given their scores on the BIS/BAS. Interestingly, the Lack of Perseverance subscale from the UPPS-P also has a significant positive loading on PC 4. (Lack of) perseverance has shown complicated and sometimes unclear results when measured against externalizing behaviors and sensitivity to consequences [1-4]. For instance, similar to the differences between PC 1 and PC 4 in the current analysis, in [2], (lack of) perseverance was significantly related to externalizing behavior measures only after controlling for the effects of reward and punishment sensitivity. Lack of Perseverance is thought to be associated with the inattentive form of ADHD [5, 6]. This fact coupled with the negative BIS/BAS loadings supports the idea that PC 4 distinguishes between inattentive (higher on PC 4) and hyperactive/impulsive (lower on PC 4) characteristics. This dynamic could help explain why females, who are more likely to suffer from inattentive symptoms [7], have higher scores on average for PC 4. PC 4 is positively related to alcohol use, indicating that perhaps CBCL scores and lack of perseverance are important predictors of early alcohol use initiation. Additionally, older participants are more likely to score high on PC 4.

Component 5 (PC 5) has significant loadings solely from SST measures of accuracy during GO trials. Higher scores on PC 5 indicate that a participant correctly responded on more GO trials and left fewer GO trials unanswered. Commonly performed analyses of SST data typically focus on reaction time patterns, as accuracy is typically controlled for with difficulty adaptation, so it might be tempting to ignore this component as unrelated to impulsivity. However, this component was positively related to alcohol use initiation, and so it may be the case that accuracy during the SST has some importance for alcohol use outcomes that should not be ignored.

PC 5 might be explained in terms of how the titration algorithm behind the GO-STOP delay interacts with responders who have a pronounced action bias [10], which is the tendency toward acting irrespective of whether the outcome is beneficial. Action bias has been documented in the go/no-go task [11] and in various decision domains outside of the laboratory [10, 12, 13]. The SST algorithm used in the ABCD study adjusts the delay between GO and STOP signals to achieve a rate of 50% accuracy on STOP trials. If, on some trials, a participant does not pay attention and produces a GO response regardless of trial type, then the titration algorithm will make the task easier until the rate of correct STOP responses equilibrates to 50%. High action bias would appear as more GO trials answered and a higher rate of correct GO responses, but there would be no change in rates of correct STOP trials due to the adaptive difficulty level. This pattern is what is found in PC5, which supports the idea that PC 5 measures a stronger-than-average bias toward action, possibly combined with a lack of effortful attention. Older participants are more likely to have high scores on PC 5, as are children with parents who attended college.

Component 6 (PC 6) has significant negative loadings from four of the five UPPS-P scales. Higher scores on PC 6 indicate that UPPS-P scores are lower than would be expected given other results. Females and older participants are more likely to have low scores on PC 6. PC 6 thus highlights the importance of considering gender and age when interpreting the UPPS-P. Young boys are more likely to have CBCL measures of externalizing symptoms that are higher than expected given their UPPS-P scores, while older girls are more likely to show the opposite relationship. PC 6 is also positively related to alcohol use initiation.

Component 7 (PC 7), like PC 5, has loadings solely from the SST. However, these loadings are from the reaction time measures rather than from the accuracy measures. Higher scores on PC 7 indicate faster reaction times on all SST trials, and particularly on GO SST trials. Alcohol use initiation is positively related to this PC. Participants who are male, older, and have higher parental education are more likely to score highly on this component. Low scores on this component may reflect the decision to strategically slow down on the task following feedback [15, 16], or they may indicate an inability to inhibit prepotent responses [17].

Component 8 (PC 8) has no significant loadings. However, there is a notable pattern among the delay discounting coefficients. PC 8 measures how steep the participant’s discounting curve is as a function of delay, with the discounting effect at one month being the same on average for all values of PC 8. High values on PC 8 correspond to relatively flat discounting curves, and low values correspond to steeper curves (see Figure S1). Notably, this variation is orthogonal to the $k$ value that is typically measured in delay discounting procedures [18]. The value of $k$, which measures how quickly subjective value falls with delay, is related to discounting rates across all temporal horizons; in contrast, PC 8 measures the difference between discounting rates for near vs. far timepoints. Female participants are more likely to have high values for PC 8, and these higher values produce flatter discounting curves. Alcohol use initiation is negatively related to this component, and therefore positively related to steeper discounting curves.

| **Table S1** |  |  |
| --- | --- | --- |
| *Sample Demographics* |  |  |
| **Category** | **Variable** | **Mean (SD)** |
| Age | Years | 9.9 (0.6) |
|  |  | **N (%)** |
| DSM Diagnosis | Conduct Disorder | 374 (3.2) |
|  | Oppositional Defiant Disorder | 1660 (14.0) |
|  | Attention-Deficit/Hyperactivity Disorder | 1261 (10.6) |
| Ethnicity | Hispanic | 2405 (20.3) |
| Race (non-Hispanic) | Asian | 251 (2.1) |
|  | Black | 1778 (15.0) |
|  | Other | 1243 (10.5) |
|  | White | 6165 (52.1) |
| Caregiver Education | < HS Diploma | 594 (5.0) |
|  | HS Diploma/GED | 1133 (9.6) |
|  | Some College | 3074 (26.0) |
|  | Bachelor | 3011 (25.4) |
|  | Post Graduate Degree | 4030 (34.0) |
| Household Yearly Income | < 50K | 3499 (25.9) |
|  | >= 50K and < 100K | 3369 (28.4) |
|  | >= 100K | 4974 (42.0) |
| Sex at Birth | Female | 5660 (47.8) |
| ABCD Site | Yale | 600 (5.1) |
|  | Washington University St. Louis | 706 (6.0) |
|  | Virginia Commonwealth University | 550 (4.6) |
|  | University of Wisconsin-Milwaukee | 384 (3.2) |
|  | University of Vermont | 576 (4.9) |
|  | University of Utah | 1009 (8.5) |
|  | University of Pittsburgh | 459 (3.9) |
|  | University of Minnesota | 607 (5.1) |
|  | University of Michigan | 724 (6.1) |
|  | University of Maryland Baltimore | 608 (5.1) |
|  | University of Florida | 453 (3.8) |
|  | UCSD | 739 (6.2) |
|  | UCLA | 432 (3.6) |
|  | SRI International | 353 (3.0) |
|  | University of Rochester | 339 (2.9) |
|  | Oregon Health and Science University | 584 (4.9) |
|  | Medical University of South Carolina | 376 (3.2) |
|  | Laureate Institute | 747 (6.3) |
|  | Florida International | 631 (5.3) |
|  | Colorado Boulder | 559 (4.7) |
|  | Children's Hospital Los Angeles | 406 (3.4) |
| Caregiver Marital Status | Married | 7967 (67.3) |
|  | Separated | 464 (3.9) |
|  | Never Married | 1454 (12.3) |
|  | Living with Partner | 668 (5.8) |
|  | Divorced | 1078 (9.1) |
|  | Widowed | 97 (0.8) |
|  | Other/Refused | 94 (0.8) |
| Caregiver Place of Birth | Outside US | 3796 (32.1) |

Table S1. Sample Demographics. In total, 11,872 individuals from 21 sites are included in this dataset. Mean and standard deviation for age in years is given. All other variables are categorical and reported by raw count and percentage.

| **Table S2** | | | | | | | | | | |  |
| --- | --- | --- | --- | --- | --- | --- | --- | --- | --- | --- | --- |
| *Significant Principal Component Loadings* | | | | | | | | | | |  |
| Source | Item |  | Principal Component | | | | | | | |  |
|  |  | | 1 | 2 | 3 | 4 | 5 | 6 | 7 | 8 |  |
|  | Percent Explained Variance | | 13.0 | 12.4 | 10.5 | 8.0 | 6.7 | 5.3 | 5.3 | 4.1 |  |
| CBCL | ADHD t-score | | **0.20** | 0.02 | **0.35** | **0.30** | 0.19 | 0.28 | 0.05 | 0.04 |  |
|  | Attention Problems t-score | | **0.19** | 0.02 | **0.34** | **0.31** | 0.18 | 0.27 | 0.04 | 0.02 |  |
|  | Externalizing Problems t-score | | **0.18** | 0.02 | **0.31** | **0.24** | 0.15 | 0.22 | 0.07 | 0.04 |  |
| BIS/BAS | BAS – Fun Seeking | | 0.14 | 0.02 | **0.25** | **-0.38** | 0.01 | 0.04 | 0.01 | 0.08 |  |
|  | BAS – Drive | | **0.14** | 0.00 | **0.24** | **-0.33** | -0.02 | 0.09 | -0.02 | 0.02 |  |
|  | BAS – Reward Responsiveness | | 0.10 | 0.01 | **0.19** | **-0.40** | -0.03 | 0.19 | 0.01 | 0.04 |  |
|  | BIS | | 0.07 | -0.01 | 0.15 | **-0.30** | 0.01 | 0.04 | -0.03 | -0.19 |  |
| UPPS-P | Lack of Perseverance | | 0.10 | 0.02 | 0.11 | **0.27** | 0.09 | **-0.48** | 0.00 | -0.01 |  |
|  | Lack of Planning | | 0.09 | 0.03 | 0.14 | 0.19 | 0.13 | **-0.50** | 0.08 | 0.13 | 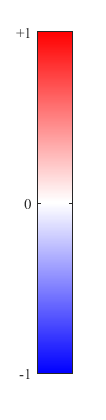 |
|  | Negative Urgency | | **0.16** | 0.01 | **0.23** | -0.17 | 0.07 | **-0.34** | 0.04 | -0.14 |  |
|  | Positive Urgency | | **0.18** | 0.00 | **0.23** | -0.17 | 0.07 | **-0.39** | -0.01 | -0.11 |  |
|  | Sensation Seeking | | 0.04 | 0.05 | 0.14 | -0.23 | 0.05 | -0.06 | 0.05 | 0.24 |  |
| SST | Correct Go Trials Mean RT | | **-0.13** | **-0.31** | 0.08 | -0.01 | 0.21 | -0.01 | **-0.48** | 0.09 |  |
|  | Correct Go Trials Count | | -0.14 | 0.02 | -0.16 | -0.13 | **0.58** | 0.03 | 0.24 | -0.15 |  |
|  | Correct Stop Trials Count | | **-0.17** | **-0.40** | 0.10 | -0.01 | 0.04 | -0.02 | 0.06 | 0.03 |  |
|  | Incorrect Go Trials Mean RT | | -0.10 | **-0.25** | 0.07 | -0.01 | 0.17 | -0.02 | **-0.53** | 0.15 |  |
|  | Incorrect Go Trials Count | | **0.18** | **0.31** | 0.01 | 0.07 | **-0.32** | 0.00 | -0.07 | 0.05 |  |
|  | Incorrect Stop Trials Mean RT | | 0.11 | **0.29** | -0.08 | 0.00 | 0.02 | -0.01 | -0.36 | 0.15 |  |
|  | Incorrect Stop Trials Count | | **0.14** | **0.43** | -0.14 | -0.02 | 0.09 | 0.02 | -0.06 | -0.03 |  |
|  | Unanswered Go Trials Count | | 0.06 | **-0.18** | 0.18 | 0.11 | **-0.54** | -0.04 | -0.12 | 0.13 |  |
|  | All Trials Mean RT | | 0.11 | **0.32** | -0.11 | -0.03 | 0.21 | 0.00 | **-0.44** | 0.07 |  |
| Delay Discounting | Six Hour Indifference Point | | **-0.27** | **0.15** | 0.11 | -0.03 | 0.03 | -0.03 | 0.11 | 0.43 |  |
|  | One Day Indifference Point | | **-0.27** | **0.15** | 0.12 | -0.04 | 0.02 | -0.04 | 0.12 | 0.42 |  |
|  | One Week Indifference Point | | **-0.32** | **0.18** | 0.16 | -0.02 | 0.00 | -0.03 | 0.05 | 0.19 |  |
|  | One Month Indifference Point | | **-0.33** | **0.18** | **0.20** | 0.02 | -0.02 | 0.00 | 0.01 | -0.01 |  |
|  | Three Month Indifference Point | | **-0.32** | **0.17** | **0.20** | 0.02 | -0.04 | -0.02 | -0.05 | -0.17 |  |
|  | One Year Indifference Point | | **-0.29** | **0.15** | **0.22** | 0.04 | -0.06 | -0.01 | -0.10 | -0.35 |  |
|  | Five Year Indifference Point | | **-0.24** | 0.13 | **0.21** | 0.05 | -0.08 | 0.01 | -0.15 | -0.46 |  |
| *Note: Cells are color-coded by the strength of their loadings. Deeper red cells have stronger positive loadings and deeper blue cells have stronger negative loadings. White cells have loadings close to 0. Bold font indicates that the loading is significant after controlling for false discoveries [1]. Abbreviations: ADHD, attention deficit/hyperactivity disorder; BIS/BAS, Behavioral Inhibition System/Behavioral Activation System; CBCL, Child Behavior Checklist; RT, reaction time; SST, stop signal task; UPPS-P, Urgency-Premeditation-Perseverance-Sensation Seeking-Positive Urgency; % EV, percent explained variance.* | | | | | | | | | | |  |

Table S2. Principal component loadings for all eight significant PCs. Cells are color-coded by the strength of their loadings, with deep blues and deep reds indicating strong negative and positive loadings, respectively. Bolded entries are significant (q < .05) after FDR correction.

**Loading Coefficient**

| **Table S3** | | | | | | | | | |
| --- | --- | --- | --- | --- | --- | --- | --- | --- | --- |
| *Demographic and Substance Use Variables Significantly Associated with PC Values* | | | | | | | | | |
|  |  | PC | | | | | | | |
| Category | Variable | 1 | 2 | 3 | 4 | 5 | 6 | 7 | 8 |
| Substance Use | Cannabis | **0.049** | -0.018 | **0.094** | -0.026 | 0.014 | 0.001 | -0.003 | -0.021 |
|  | Alcohol | **0.074** | **-0.074** | **0.097** | **0.050** | **0.045** | **0.069** | **0.037** | **-0.052** |
| Caregiver Education | Post-Graduate | **-0.106** | -0.007 | **-0.066** | 0.028 | 0.030 | -0.014 | -0.002 | -0.005 |
|  | >= Bachelor’s | **-0.125** | **-0.088** | -0.045 | -0.025 | 0.033 | 0.016 | 0.027 | **-0.042** |
|  | >= Some college | -0.012 | -0.029 | 0.045 | -0.029 | **0.045** | -0.034 | **0.057** | -0.006 |
|  | >= HS Diploma | -0.020 | -0.018 | 0.003 | 0.019 | 0.012 | 0.009 | **0.041** | -0.022 |
| Household Yearly Income | > $100,000/yr | -0.027 | 0.011 | -0.035 | 0.018 | -0.003 | -0.011 | 0.018 | -0.009 |
|  | > $50,000/yr | **-0.072** | -0.035 | -0.043 | 0.019 | -0.016 | 0.014 | 0.009 | 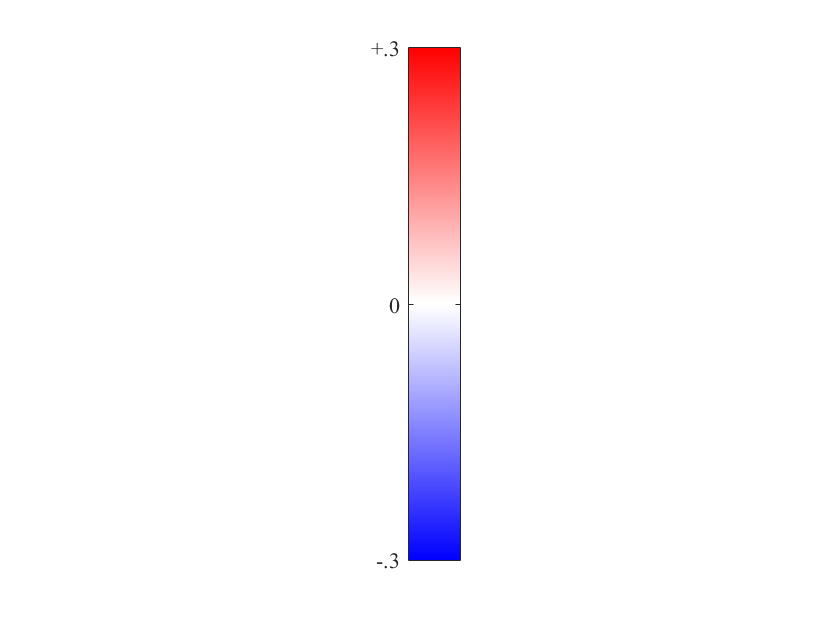-0.022  **Regression Coefficient** |
| Sex at Birth | Female | **-0.281** | **0.222** | **-0.074** | -0.008 | -0.004 | **-0.065** | **-0.165** | **0.103** |
| Age | Years | **-0.068** | **-0.057** | **-0.098** | **0.034** | **0.043** | **-0.050** | **0.278** | 0.005 |
| ABCD Site | Yale | -0.038 | **0.115** | **0.078** | 0.012 | **0.089** | 0.024 | **-0.079** | **-0.051** |
|  | Washington University St. Louis | 0.036 | -0.021 | 0.055 | 0.014 | 0.025 | 0.041 | **-0.048** | -0.022 |
|  | Virginia Commonwealth University | -0.042 | **0.118** | 0.017 | -0.046 | 0.038 | **0.050** | -0.037 | 0.011 |
|  | University of Wisconsin-Milwaukee | -0.013 | 0.040 | 0.052 | -0.027 | 0.031 | 0.019 | 0.013 | -0.002 |
|  | University of Vermont | -0.019 | **0.106** | **0.100** | -0.040 | -0.011 | **0.043** | -0.005 | **-0.049** |
|  | University of Utah | **-0.077** | 0.051 | **0.157** | -0.011 | **0.135** | -0.022 | 0.019 | -0.008 |
|  | University of Pittsburgh | 0.056 | **0.067** | **0.121** | -0.025 | 0.030 | 0.008 | **-0.056** | -0.015 |
|  | University of Minnesota | -0.064 | **0.094** | -0.006 | -0.043 | -0.002 | **0.054** | 0.018 | -0.010 |
|  | University of Michigan | -0.024 | 0.066 | 0.060 | -0.010 | **0.073** | 0.010 | **-0.183** | **-0.079** |
|  | University of Maryland Baltimore | 0.005 | 0.059 | **0.112** | **-0.054** | 0.013 | -0.005 | 0.033 | 0.000 |
|  | University of Florida | 0.025 | **0.061** | **0.110** | 0.017 | -0.020 | -0.010 | 0.026 | -0.017 |
|  | UCSD | -0.017 | 0.046 | **0.062** | 0.050 | **0.066** | 0.015 | -0.018 | -0.029 |
|  | UCLA | -0.010 | 0.034 | 0.037 | 0.001 | 0.040 | -0.022 | **-0.111** | **-0.059** |
|  | SRI International | -0.020 | **0.063** | **0.077** | -0.015 | **0.159** | 0.003 | **-0.245** | **-0.091** |
|  | University of Rochester | 0.039 | -0.036 | **0.085** | -0.002 | 0.018 | 0.031 | 0.020 | -0.014 |
|  | Oregon Health and Science University | -0.056 | 0.037 | **0.068** | -0.052 | **0.101** | 0.030 | **-0.105** | **-0.071** |
|  | Medical University of South Carolina | -0.017 | **0.132** | **0.088** | 0.035 | **0.055** | **0.050** | **-0.110** | **-0.050** |
|  | Laureate Institute | 0.011 | **0.115** | **0.108** | -0.007 | **0.090** | -0.034 | **-0.105** | -0.036 |
|  | Florida International | -0.021 | **0.099** | **0.126** | 0.002 | **0.070** | **-0.080** | **-0.068** | -0.026 |
|  | Colorado Boulder | -0.053 | **0.122** | 0.036 | 0.041 | **0.100** | 0.008 | **-0.274** | **-0.107** |
| Caregiver Marital Status | Married | -0.040 | -0.088 | 0.045 | 0.095 | 0.036 | 0.022 | -0.028 | -0.008 |
|  | Separated | 0.037 | -0.056 | 0.045 | 0.010 | 0.006 | -0.038 | 0.001 | -0.006 |
|  | Never Married | 0.077 | -0.051 | 0.108 | 0.010 | 0.014 | -0.031 | -0.030 | -0.012 |
|  | Living with Partner | 0.035 | -0.036 | 0.060 | 0.042 | 0.009 | -0.020 | 0.001 | -0.004 |
|  | Divorced | 0.016 | -0.046 | 0.044 | 0.029 | 0.021 | 0.009 | -0.021 | -0.004 |
|  | Widowed | 0.017 | -0.048 | 0.021 | -0.015 | 0.019 | 0.009 | 0.003 | 0.006 |
| *Note: Regression coefficients are standardized. Cells are color-coded by the strength of their regression coefficients. Deeper red cells have stronger positive coefficients and deeper blue cells have stronger negative coefficients. White cells have loadings close to 0. Bold font indicates that the coefficient is significant after controlling for false discoveries [1]. Besides age, all variables are binary.* | | | | | | | | | |

Table S3. Significant associations with PCs 1-8. Entries are standardized regression coefficients where the PC value is the dependent variable and all entries in the table are independent variables. Binary variables, site, and marital status values are dummy coded. Household yearly income and caregiver education level are ordinal coded. Cells are color coded by the strength of their regression coefficients, with deep blues and deep reds indicating strong negative and positive coefficients. Bolded entries are significant (q < .05) after FDR correction.

PC 8 Average Delay Discounting Curves

Current value of $100

| Delay in log(seconds) | mean | 1 SD above | 1 SD below |
| --- | --- | --- | --- |
| 0.778151 | 85.47 | 74.1602 | 96.7798 |
| 1.38021 | 83.86 | 72.4944 | 95.2256 |
| 2.22531 | 74.91 | 68.8425 | 80.9775 |
| 2.85733 | 67.05 | 67.5347 | 66.5653 |
| 3.33445 | 60.27 | 66.321 | 54.219 |
| 3.9425 | 50.31 | 63.449 | 37.171 |
| 4.64147 | 38.27 | 55.5764 | 20.9636 |

Figure S1. Average discounting curves for three values of PC 8. The curves correspond to the average discounting values at each time interval for individuals who have scores of 0, 1, and -1 on PC 8. The x-axis is displayed on a logarithmic scale, with time values ranging from 6 hours to 5 years. The y-axis indicates the current valuation of $100 at a specified delay of receipt measured logarithmically. As the value of PC 8 increases, the discounting curve flattens.

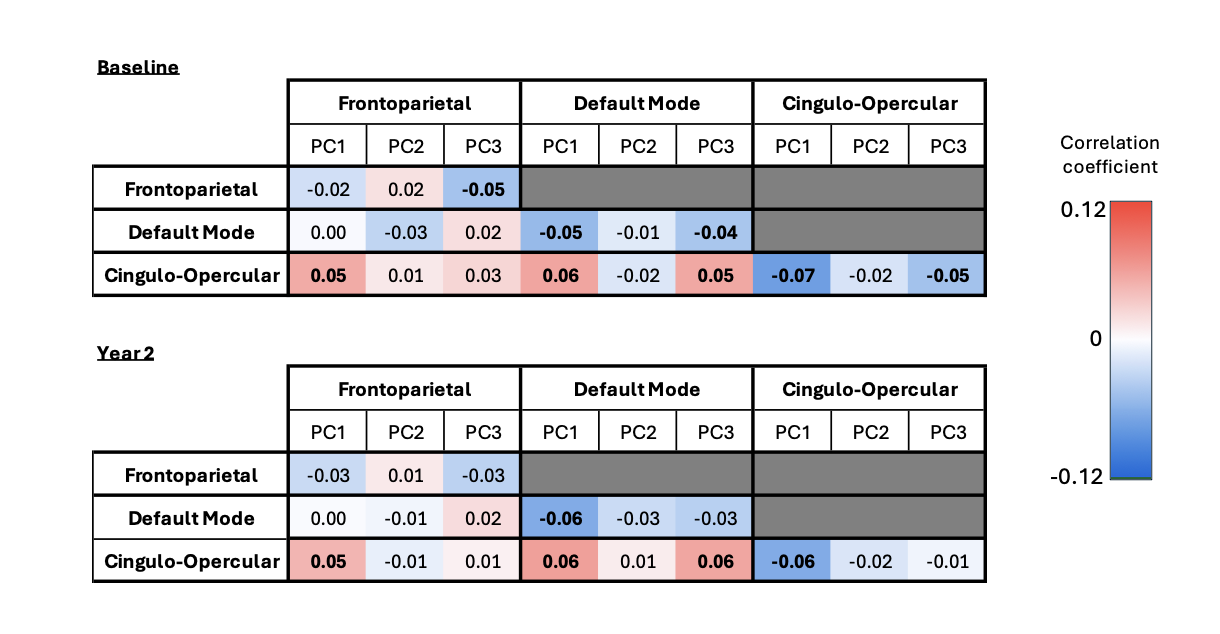

Figure S2. Partial correlation analysis controlling for mean motion for the associations between the top three principal components and all possible combinations of connectivity between and within default mode, frontoparietal, and cingulo-opercular networks at baseline and at 2-year follow-up assessments. Bold font indicates that the coefficient is significant after controlling for false discovery rate. We observed a similar pattern of association as those reported in the main manuscript.

References

1.	Maneiro, L., et al., *Impulsivity traits as correlates of antisocial behaviour in adolescents.* Personality and individual differences, 2017. **104**: p. 417-422.

2.	Carlson, S.R., A.A. Pritchard, and R.M. Dominelli, *Externalizing behavior, the UPPS-P impulsive behavior scale and reward and punishment sensitivity.* Personality and individual differences, 2013. **54**(2): p. 202-207.

3.	Whiteside, S.P. and D.R. Lynam, *The five factor model and impulsivity: Using a structural model of personality to understand impulsivity.* Personality and individual differences, 2001. **30**(4): p. 669-689.

4.	Hecht, L.K. and R.D. Latzman, *Revealing the nuanced associations between facets of trait impulsivity and reactive and proactive aggression.* Personality and Individual Differences, 2015. **83**: p. 192-197.

5.	Gomez, R. and S. Watson, *Associations of UPPS-P negative urgency and positive urgency with ADHD dimensions: Moderation by lack of premeditation and lack of perseverance in men and women.* Personality and Individual Differences, 2023. **206**: p. 112125.

6.	Lopez, R., et al., *A multidimensional approach of impulsivity in adult attention deficit hyperactivity disorder.* Psychiatry research, 2015. **227**(2-3): p. 290-295.

7.	Nussbaum, N.L., *ADHD and female specific concerns: a review of the literature and clinical implications.* Journal of attention disorders, 2012. **16**(2): p. 87-100.

8.	Coxe, S., M.H. Sibley, and S.P. Becker, *Presenting problem profiles for adolescents with ADHD: differences by sex, age, race, and family adversity.* Child and Adolescent Mental Health, 2021. **26**(3): p. 228-237.

9.	Scheres, A., et al., *Temporal and probabilistic discounting of rewards in children and adolescents: effects of age and ADHD symptoms.* Neuropsychologia, 2006. **44**(11): p. 2092-2103.

10.	Patt, A. and R. Zeckhauser, *Action bias and environmental decisions.* Journal of Risk and Uncertainty, 2000. **21**: p. 45-72.

11.	Guitart-Masip, M., et al., *Go and no-go learning in reward and punishment: interactions between affect and effect.* Neuroimage, 2012. **62**(1): p. 154-166.

12.	Bar-Eli, M., et al., *Action bias among elite soccer goalkeepers: The case of penalty kicks.* Journal of economic psychology, 2007. **28**(5): p. 606-621.

13.	Kiderman, A., et al., *in primary care: Evidence of action bias.* The Journal of Family Practice, 2013. **62**(8).

14.	Logan, G.D. and W.B. Cowan, *On the ability to inhibit thought and action: A theory of an act of control.* Psychological review, 1984. **91**(3): p. 295.

15.	Bissett, P.G. and G.D. Logan, *Post-stop-signal slowing: strategies dominate reflexes and implicit learning.* Journal of Experimental Psychology: Human Perception and Performance, 2012. **38**(3): p. 746.

16.	Boehler, C.N., et al., *The influence of different Stop-signal response time estimation procedures on behavior–behavior and brain–behavior correlations.* Behavioural brain research, 2012. **229**(1): p. 123-130.

17.	Aron, A.R., *The neural basis of inhibition in cognitive control.* The neuroscientist, 2007. **13**(3): p. 214-228.

18.	Odum, A.L., *Delay discounting: I'm ak, you're ak.* Journal of the experimental analysis of behavior, 2011. **96**(3): p. 427-439.
